# Fatty Acid Pathways Regulate Thermal Nociception in *Caenorhabditis elegans*

**DOI:** 10.1101/2025.10.17.683074

**Authors:** Marzieh Abdollahi, Francis Beaudry

**Author notes:** Corresponding author: Francis Beaudry, Ph.D., Professor of Analytical Pharmacology, Canada Research Chair in metrology of bioactive molecules and target discovery, Département de Biomédecine Vétérinaire, Faculté de Médecine Vétérinaire, Université de Montréal, 3200 Sicotte, Saint-Hyacinthe, QC, Canada J2S 2M2.

## Abstract

Chronic pain remains a major unmet medical challenge, and lipid signaling pathways have emerged as key modulators of nociception. Using *Caenorhabditis elegans* as a genetically tractable model, we investigated how fatty acid composition influences thermal avoidance behavior. Mutant strains lacking functional desaturase enzymes (*elo-1, fat-1, fat-2, fat-3, fat-4, fat-6*/*fat-7*), and consequently depleted in polyunsaturated fatty acids (PUFAs) such as arachidonic acid, displayed significantly reduced sensitivity to noxious heat compared to wild-type animals. These findings indicate that intact PUFA biosynthesis is essential for normal thermal nociception in *C. elegans*. Given that arachidonic acid is a precursor of endocannabinoids (AEA and 2-AG) known to modulate TRPV1-dependent pain signaling, our results suggest that a conserved lipid-based mechanism regulates heat avoidance in nematodes. This study establishes a functional link between fatty acid metabolism and nociceptive behavior, providing a powerful platform to explore metabolic modulation of pain pathways.

## Introduction

Arachidonic acid (AA), a polyunsaturated omega-6 fatty acid, is a pivotal mediator linking metabolism, inflammation, and pain [1–3]. In mammals, AA is synthesized from dietary linoleic acid (LA) through sequential desaturation and elongation by Δ6- and Δ5-desaturases, yielding γ-linolenic acid (GLA) and dihomo-γ-linolenic acid (DGLA) as intermediates [4–6]. This pathway provides the substrate for eicosanoids, potent lipid mediators that orchestrate inflammatory and nociceptive signaling. Upon cellular stimulation or injury, phospholipase A2 (PLA2) liberates AA from membrane phospholipids. Free AA is then metabolized by cyclooxygenases (COX) and lipoxygenases (LOX) into prostaglandins, thromboxanes, and leukotrienes [6]. Prostaglandins, in particular, sensitize nociceptors and drive vasodilation, edema, and fever, thereby amplifying pain transmission [7–9].

Beyond its role in eicosanoid signaling, AA is the biochemical foundation of the endocannabinoid system, a lipid signaling network that regulates pain, mood, and immune function [10,11]. The principal endocannabinoids, anandamide (AEA) and 2-arachidonoylglycerol (2-AG), are synthesized from AA-containing phospholipids via NAPE-specific phospholipase D (NAPE-PLD) and diacylglycerol lipase (DAGL), respectively. Once released, AEA and 2-AG activate cannabinoid receptors CB1 and CB2 to suppress nociceptive transmission. Their signaling is rapidly terminated by fatty acid amide hydrolase (FAAH) and monoacylglycerol lipase (MAGL), ensuring tight temporal control [7]. The endocannabinoid system thus acts as a critical homeostatic modulator of pain [13–19]. CB1/CB2 activation inhibits neurotransmitter release, while AEA and 2-AG also engage the TRPV1 channel, integrating cannabinoid and nociceptive pathways [8,9]. Endocannabinoids further suppress pro-inflammatory cytokines and enhance anti-inflammatory responses, attenuating both inflammatory and neuropathic pain [^1–5^]. Together, these mechanisms position the endocannabinoid system as a central regulator of pain processing and a compelling target for next-generation analgesic therapies.

Despite extensive research on endocannabinoid signaling in mammals, its evolutionary origins and function in invertebrates remain poorly understood [10–12]. The discovery of endocannabinoid molecules such as anandamide (AEA) and 2-arachidonoylglycerol (2-AG), along with the identification of cannabinoid-like receptors NPR-19 and NPR-32 in *Caenorhabditis elegans* (*C. elegans*), has revealed a conserved molecular framework for endocannabinoid-like signaling [10,11]. These findings suggest that *C. elegans* harbors the essential components of a functional lipid signaling system capable of modulating nociceptive responses. Arachidonic acid (AA) availability likely constrains the synthesis of these signaling mediators. In *C. elegans*, AA is produced through coordinated activity of desaturase (FAT) and elongase (ELO) enzymes (Figure 2) [6]. Together, FAT and ELO catalyze the conversion of dietary fatty acids into long-chain polyunsaturated products, including AA, a critical precursor for prostaglandins, leukotrienes, and endocannabinoid-like lipids that regulate inflammation and nociception. Insufficient AA biosynthesis may thus impair the organism’s ability to mount appropriate responses to noxious stimuli. Leveraging *C. elegans* as a genetically tractable model provides a unique opportunity to dissect the molecular mechanisms and evolutionary basis of endocannabinoid signaling. We hypothesize that *fat* and *elo* mutants, deficient in AA biosynthesis, will exhibit hindered avoidance of noxious heat due to disrupted production of lipid mediators. Such phenotypes would highlight the pivotal role of fatty acid metabolism in modulating nociceptive and thermosensory pathways. By characterizing these mutants, we aim to establish a mechanistic link between lipid metabolism, endocannabinoid-like signaling, and behavioral responses to pain-related stimuli.

**Figure 1.**
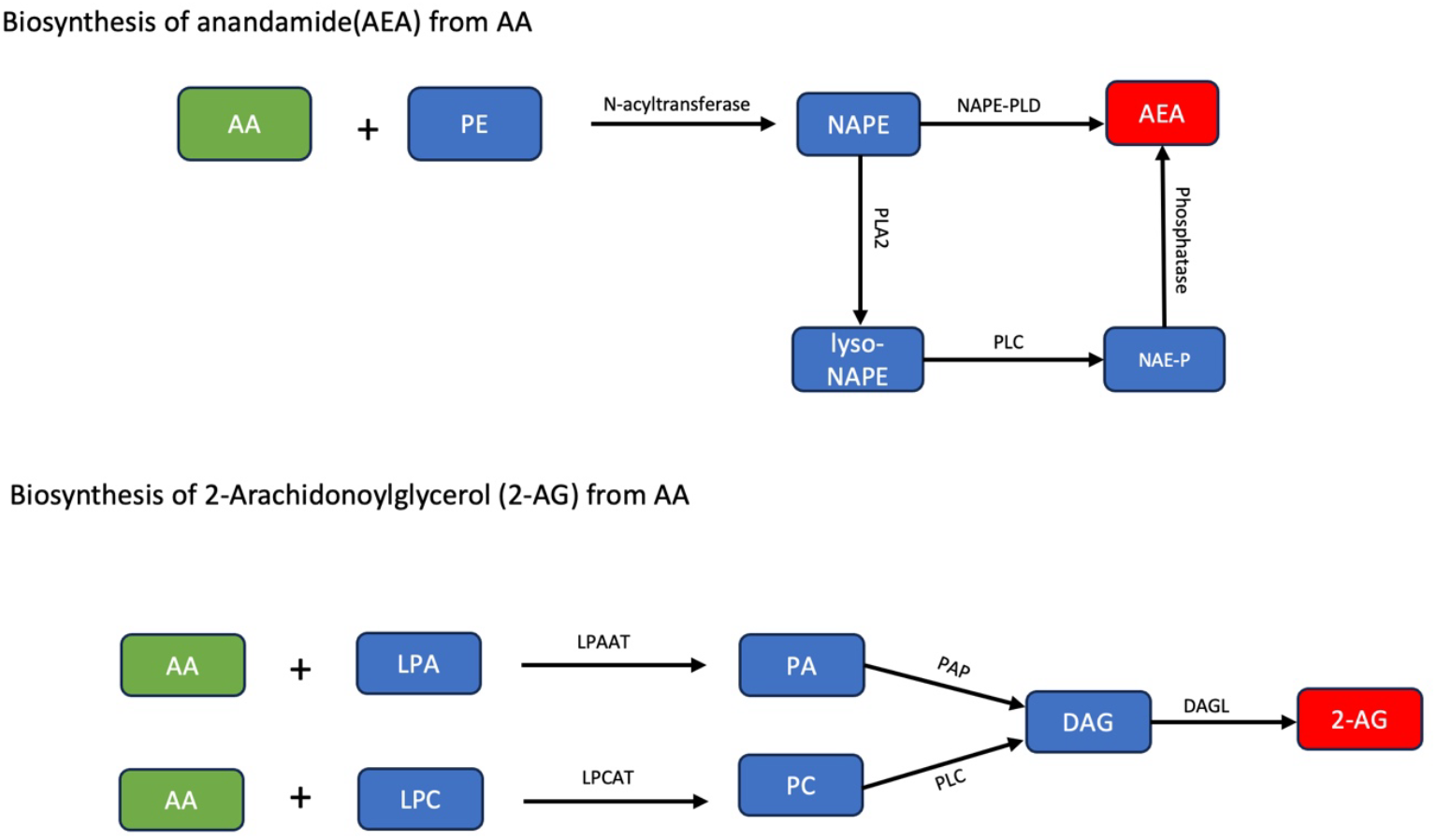
Biosynthesis of AEA and 2-AG from AA

**Figure 2.**
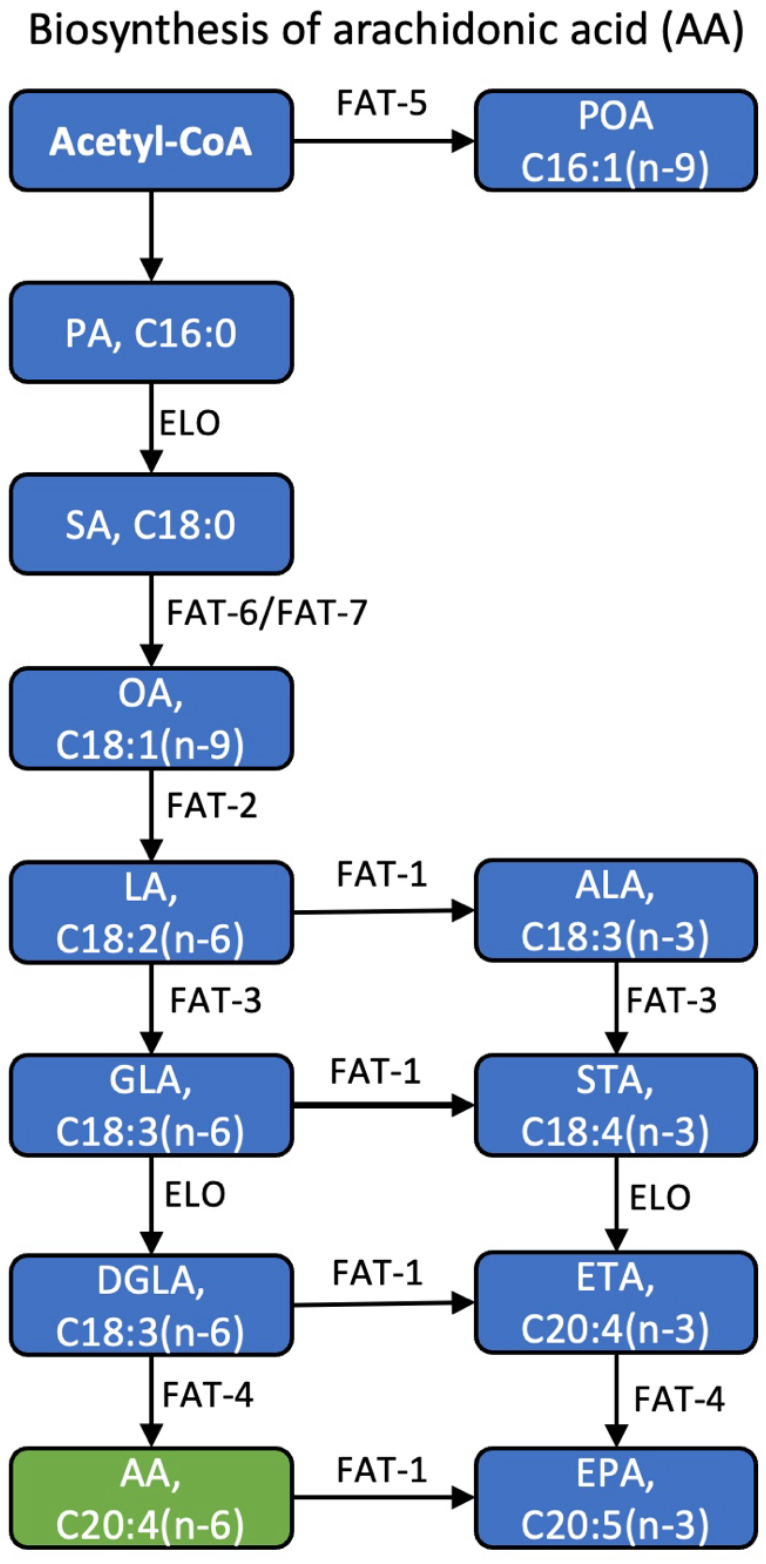
Biosynthesis of AA in *C. elegans* and the role of FAT and ELO enzymes

## Materials and methods

### Chemicals and Reagents

All chemicals and reagents were obtained from Fisher Scientific (Fair Lawn, NJ, USA) or Millipore Sigma (St. Louis, MO, USA), unless otherwise specified. Capsaicin (Cap) was purchased from Toronto Research Chemicals (North York, ON, Canada), Δ^9^-tetrahydrocannabinol (THC) from Cerilliant (Round Rock, TX, USA), and anandamide (AEA) from Cayman Chemical (Ann Arbor, MI, USA).

### *C. elegans* Strains and Maintenance

The *Caenorhabditis elegans* N2 (Bristol) strain was used as the wild-type reference. The following mutant strains were included: *elo-1* (BX14), *fat-6;fat-7* (BX156), *fat-2* (BX26), *fat-3* (BX30), *fat-4* (BX17), *fat-1* (BX24), and *fat-4;fat-1* (BX52). All strains were obtained from the Caenorhabditis Genetics Center (CGC), University of Minnesota (Minneapolis, MN, USA). Nematodes were cultured and handled under standard conditions as previously described [13]. Worms were maintained on nematode growth medium (NGM) agar plates seeded with *Escherichia coli* OP50 at 22 °C in a refrigerated incubator (Heratherm, Thermo Scientific, Fair Lawn, NJ, USA). Unless otherwise indicated, all behavioral and biochemical assays were performed at room temperature (∼22 °C).

## Thermal avoidance assays

The behavioral analysis technique for assessing thermal avoidance was adapted from the previously detailed four-quadrant strategy ^6^. This method has been extensively applied in our earlier studies ^7–13^. In summary, the experiments utilized 92 × 16 mm Petri dishes segmented into four quadrants. The nematodes used during the experiments were off-food. A central circle (1 cm in diameter) marked an area where *C. elegans* was not considered. The quadrants of the Petri dishes were designated as two stimulus areas (A and D) and two control areas (B and C). Sodium azide (0.5 M) was employed to immobilize the nematodes in all quadrants. A radial temperature gradient (32–35°C on NGM agar 2 mm from the tip) was created with an electronically heated metal tip (0.8 mm in diameter), and the temperature was measured using an infrared thermometer. The stimulus temperature was based on prior experiments ^14^. Nematodes were isolated and cleaned following the protocol by Margie et al. ^6^. Typically, 100 to 300 young adult nematodes were placed at the center of a marked Petri dish, and after 30 minutes, the plates were kept at 4°C for at least 1 hour before counting the number of nematodes per quadrant. Nematodes that did not cross the inner circle were excluded. The thermal avoidance index (TI) was calculated using Equation 1.

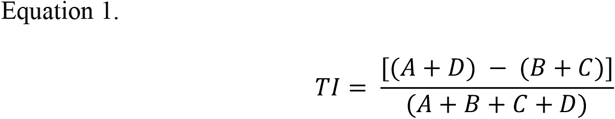

The technical details are presented in Supplementary Figure S1. Both TI and animal avoidance percentages were used to analyze nocifensive responses to noxious heat in each tested *C. elegans* experimental group.

## Statistical analysis

Behavioral data were analyzed using the Kruskal-Wallis test, a non-parametric method, followed by the post-hoc Dunn test for multiple comparisons. Statistical significance was predetermined at p ≤ 0.05. All analyses were conducted using GraphPad PRISM (version 10.6.0).

## Results and discussion

In a recent study, we demonstrated that AEA modulates thermal avoidance in *C. elegans* through the interplay between vanilloid and cannabinoid receptors [14]. Our findings suggest that AEA, exerts its effects by engaging both receptor types, which work together to trigger the organism’s response to noxious heat. This discovery underscores the complex signaling mechanisms underlying thermal nociception and highlights the role of endocannabinoid signaling in regulating behavioral responses to environmental stressors. Impaired AA production in *C. elegans*, due to dysfunctional FAT and ELO enzymes, could lead to reduced AEA levels. This deficiency may disrupt AEA signaling through vanilloid and cannabinoid receptors, impairing the nematodes’ ability to effectively avoid noxious heat.

In mammals, AEA modulates pain perception, reduces inflammation, and inhibits pain signals ^15–18^. The first step was to determine whether any behavioral bias existed in *C. elegans* under our experimental conditions. To assess this, we evaluated the mobility and quadrant preference of both wild-type (N2) and specific mutant nematodes. The experiments were conducted at room temperature (≈ 22°C), ensuring consistency across all quadrants. As shown in Fig. 3, our results indicate no significant quadrant selection bias among any of the *C. elegans* experimental groups. The data demonstrate a uniform distribution of nematodes across the Petri dish 30 minutes after being placed at the center. Consequently, no experimental bias was observed.

**Figure 3.**
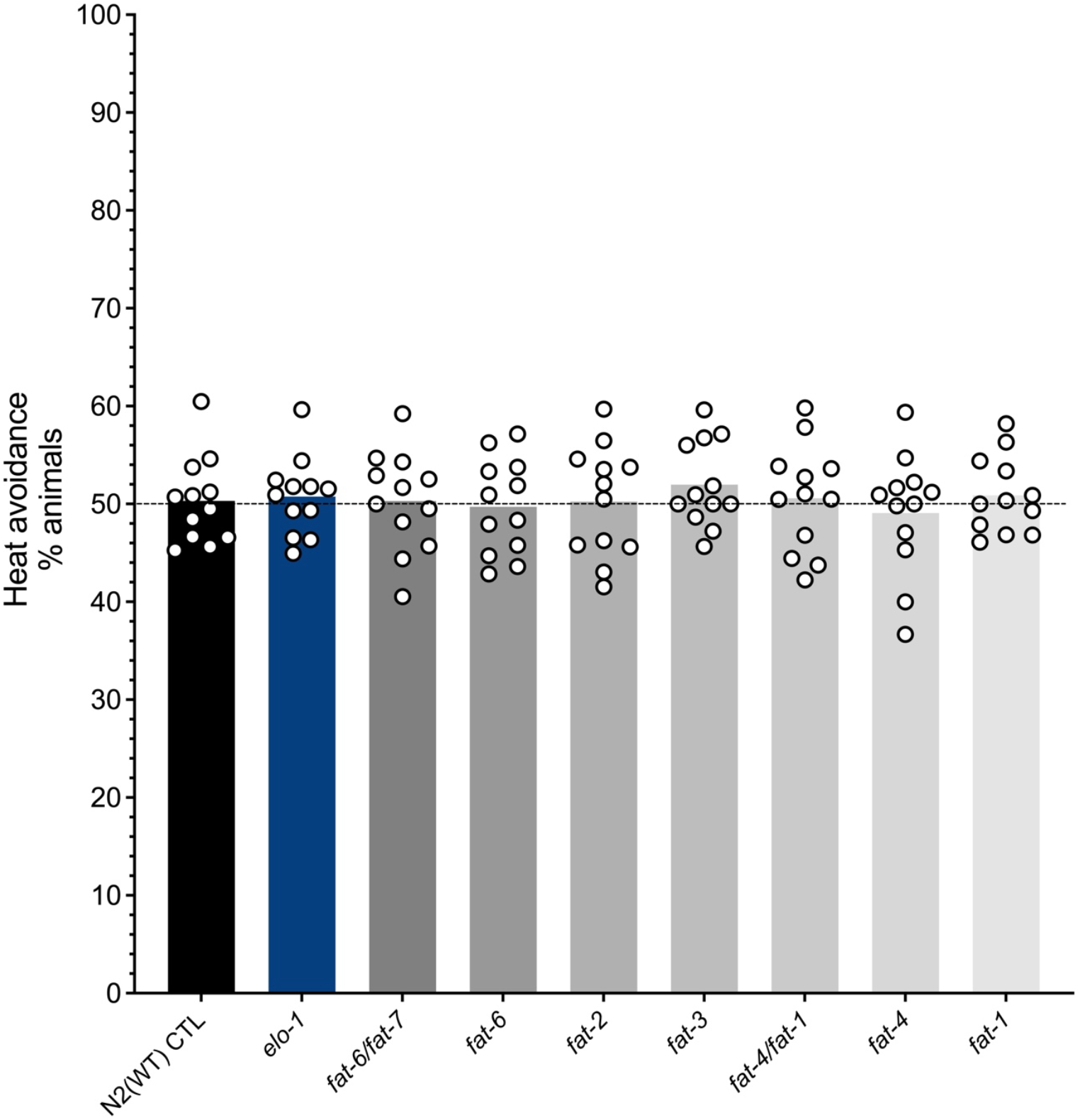
Comparison of the mobility and bias of WT (N2) and selected mutant nematodes in plates divided into quadrants conserved at a constant temperature (22°C) without the application of a stimulus (negative control). No quadrant selection bias was observed for any *C. elegans* genotype tested

Next, we investigated the heat avoidance phenotype in specific *fat* and *elo* mutants following exposure to noxious heat. As shown in Fig. 4, wild-type (N2) nematodes effectively avoid noxious heat (i.e. 32-35°C). In contrast, all tested mutants, including *elo- 1, fat-1, fat-2, fat-3, fat-4*, and double mutants *fat-6*/*fat-7* and *fat-4*/*fat-1*, exhibited significant impairments in heat avoidance behavior (i.e. 32-35°C). Our results demonstrate that perturbations in fatty acid biosynthesis profoundly alter thermal nociception in *C. elegans*. Mutants deficient in key desaturase enzymes, including elo-1, fat-1, fat-2, fat-3, fat-4, and fat-6/fat-7, exhibited a marked reduction in thermal avoidance behavior when exposed to noxious heat. This consistent phenotype across multiple genotypes implicates polyunsaturated fatty acids (PUFAs) and their downstream metabolites as critical modulators of heat sensitivity and nocifensive responses. PUFAs are precursors for bioactive lipids such as arachidonic acid (AA), derived endocannabinoids, including anandamide (AEA) and 2-arachidonoylglycerol (2-AG), both of which modulate nociceptive signaling in mammals. The attenuation of heat avoidance in PUFA-deficient mutants suggests that *C. elegans* employs a conserved lipid signaling mechanism to regulate thermal nociception. Previous studies have shown that AA and its derivatives can modulate transient receptor potential (TRP) channels, particularly TRPV homologs, thereby influencing neuronal excitability ^15,18,19^. Given that the *C. elegans* homologs of mammalian TRPV1, such as OSM-9 and OCR-2, play essential roles in thermosensation and aversive behavior ^20^, the loss of PUFA-derived signaling molecules likely dampens TRP-mediated neuronal activation in response to noxious heat. Among the mutants tested, *fat-6*/*fat-7* nematodes showed the strongest impairment in heat avoidance, consistent with their upstream role in the synthesis of long-chain and monounsaturated fatty acids. This suggests that early disruptions in the desaturation–elongation cascade exert cumulative effects on the biosynthesis of AA and other signaling lipids. Interestingly, *fat-1* mutants, which lack the ω-3 desaturase necessary for eicosapentaenoic acid production, also displayed reduced avoidance, supporting a broader contribution of both ω-6 and ω-3 PUFA pathways in maintaining nociceptive competence.

**Figure 4.**
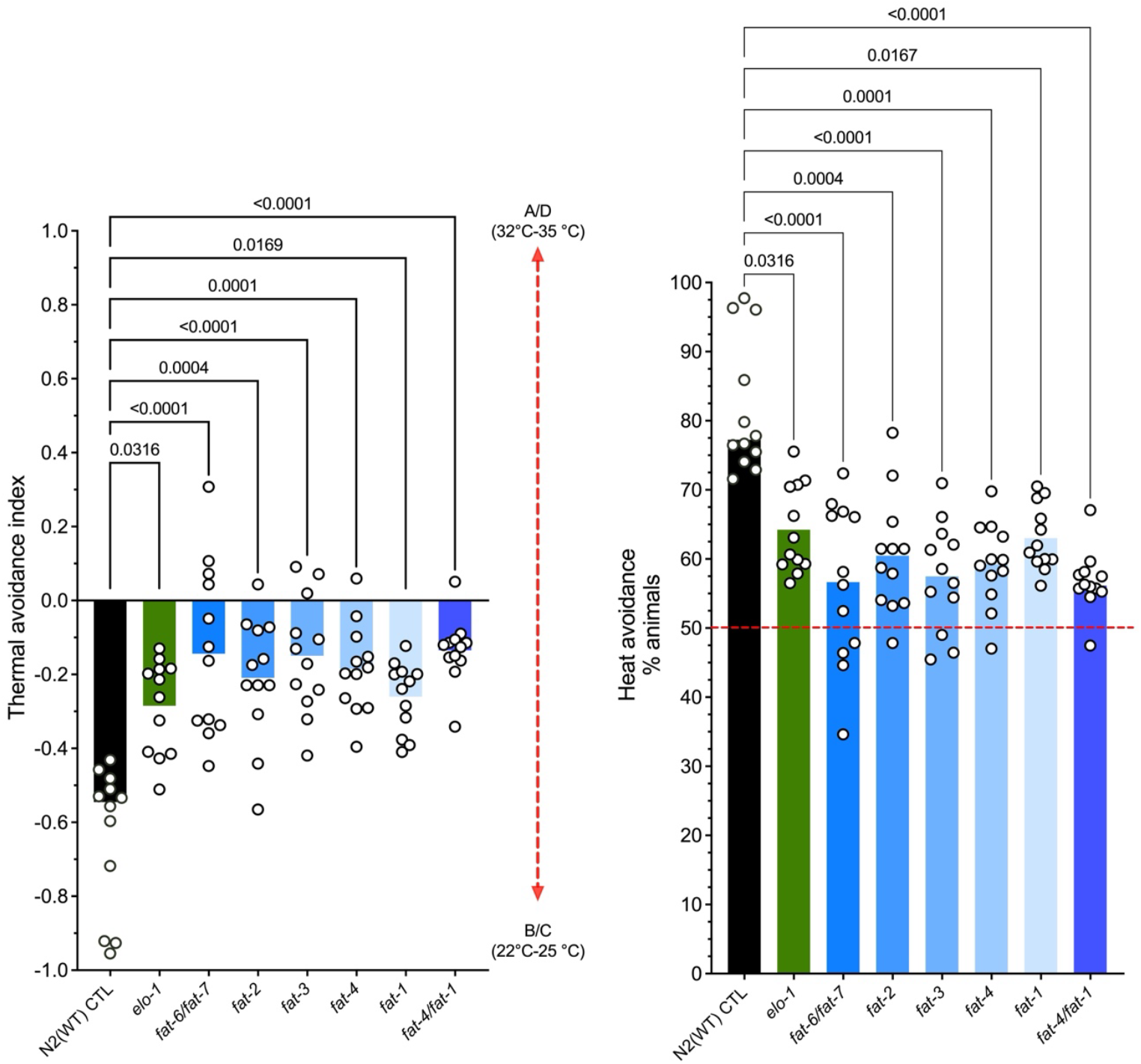
Fatty acid composition modulates thermal avoidance in C. elegans. Mutant strains lacking functional desaturase enzymes are specifically depleted in certain polyunsaturated fatty acids, including arachidonic acid, a precursor of AEA and 2-AG. Individual data points and medians are shown, derived from at least 12 independent experiments per group. All mutants tested, which are involved in the fatty acid biosynthesis pathway, exhibited significantly reduced sensitivity to noxious heat.

These findings extend the understanding of lipid-mediated nociception beyond mammalian models and establish *C. elegans* as a genetically tractable system to dissect the interplay between lipid metabolism and sensory signaling. The conservation of endocannabinoid-like pathways in nematodes underscores the evolutionary importance of lipid signaling in nociception and offers a valuable framework to explore how metabolic disruptions influence sensory homeostasis. Future work should aim to quantify the specific PUFA and endocannabinoid derivatives in these mutants and determine their direct effects on TRPV homolog activity using electrophysiological or calcium imaging approaches. Additionally, rescuing the thermal avoidance phenotype through dietary PUFA supplementation or pharmacological modulation of lipid signaling could provide causal evidence linking fatty acid metabolism to nociceptive behavior. Collectively, our data reveal that fatty acid composition is a key determinant of heat-avoidance behavior in *C. elegans*, highlighting an ancient and conserved biochemical axis between lipid metabolism and nociceptive function.

## Acknowledgement

The Université de Montréal partially provided partial financial support to M. Abdollahi. This project was funded by the National Sciences and Engineering Research Council of Canada (F. Beaudry discovery grant no. RGPIN-2020-05228). F. Beaudry is the holder of the Canada Research Chair in metrology of bioactive molecules and target discovery (grant no. CRC-2021-00160). This research was undertaken, partly, thanks to funding from the Canada Research Chairs Program.

