## Supplemental Figure 1 for "Fatty Acid Pathways Regulate Thermal Nociception in *Caenorhabditis elegans*"

**Figure S1.** A schematic of the quadrants assay adapted from Margie *et al*. (2013). For head avoidance assay, plates were divided into quadrants two test (A and D) and two controls (B and C). Sodium azide was added to all four quadrants to paralyze nematodes. *C. elegans* were added at the center of the plate (typically, n = 100 to 1,000) and after 30 minutes, animals were counted on each quadrant. Only animals outside the inner circle were scored. The calculation of thermal avoidance index was performed has described.

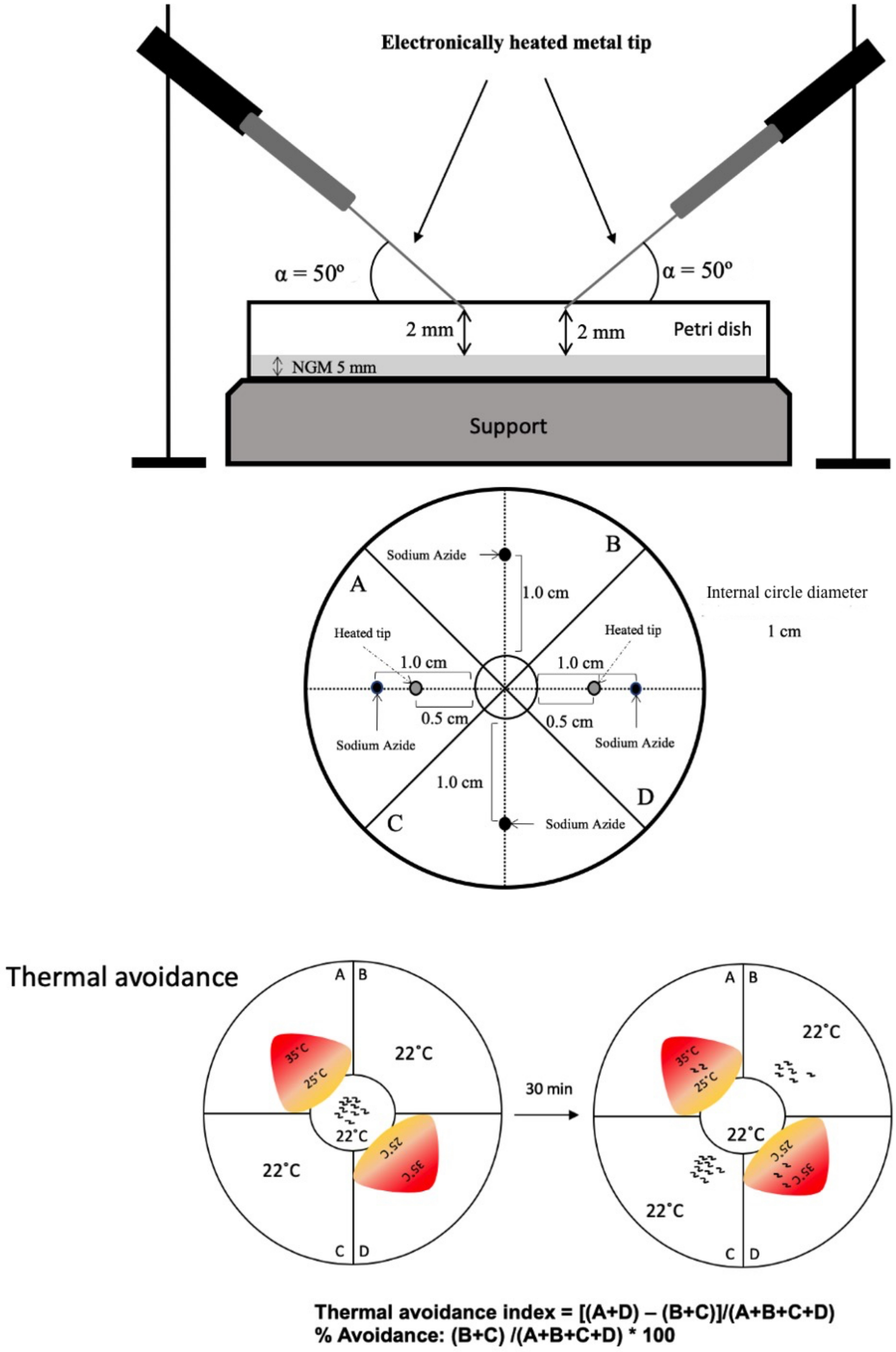
